# Distinct matrix composition and mechanics in aged and estrogen-deficient mouse skin

**DOI:** 10.1101/570481

**Authors:** Charis R Saville, Venkatesh Mallikarjun, David F Holmes, Elaine Emmerson, Brian Derby, Joe Swift, Michael J Sherratt, Matthew J Hardman

## Abstract

Hormone deficiency has been widely linked to accelerated tissue ageing, and increased incidence of chronic degenerative disease. Furthermore, age-associated hormonal dysregulation is thought to be a major contributing factor to the increased fragility of aged skin. The ageing process is driven by an aggregation of damage to cells and extracellular matrix, which can directly influence the mechanical properties of the tissue. Here we report on the correlation between mechanical properties and composition of skin from ovariectomised and aged mice, to assess the extent to which estrogen deprivation drives dermal ageing. We found that age and estrogen abrogation affected skin mechanical properties in contrasting ways: ageing led to increased tensile strength and stiffness while estrogen deprivation had the opposite effect. Mass spectrometry proteomics showed that the quantity of extractable fibrillar collagen-I decreased with ageing, but no change was observed in ovariectomised mice. This observation, in combination with measurements of tensile strength, was interpreted to reflect changes to the extent of extracellular matrix crosslinking, supported by a significant increase in the staining of advanced glycation endpoints in aged skin. Loss of mechanical strength in the skin following ovariectomy was consistent with a loss of elastic fibres. Other changes in extracellular matrix composition broadly correlated between aged and ovariectomised mice, confirming the important role of estrogen-related pathways in ageing. This study offers new insight into the relationship between tissue composition and mechanics, and suggests that the deleterious effects of intrinsic skin ageing are compounded by factors beyond hormonal dysregulation.

## INTRODUCTION

Biological ageing is associated with tissue atrophy (Fenske and Lober 1986), chronic low level inflammation (Franceschi and Campisi 2014) and a consequent impairment of normal function. In humans, intrinsically aged skin (i.e. not exposed to damaging radiation) is characterised by flattening of the dermal epidermal junction, thinning of the epidermis and loss of extracellular matrix (ECM) proteins from the dermis. Remodelling of the dermal ECM is thought to cause alterations to the mechanical properties of skin (Daly and Odland 1979), increasing the stiffness but reducing resilience and tensile strength. As a consequence, aged skin is more fragile and susceptible to injury, which in addition to other compounding age-associated factors, leads to a much higher incidence of chronic wounds in the elderly (Sgonc and Gruber 2013), incurring a healthcare burden of up to £5.1bn annually in the United Kingdom (Guest, et al. 2015).

Changes in hormonal signalling pathways represent a prominent, evolutionarily conserved feature of ageing (Broue, et al. 2007). Following menopause, women report a rapid onset of multiple signs of skin ageing, including thinning, fine wrinkles, dryness, hyperpigmentation and fragility (Brincat 2000) – concomitant with hormonal changes including a decrease in estrogen and androgen levels (reviewed in Al-Azzawi and Palacios 2009). There is good evidence that these effects can be reversed by hormone replacement therapy (HRT), which restores hydration, thickness and reduces wrinkles (Phillips, et al. 2001; Sator, et al. 2001). The mechanism for these beneficial estrogenic effects may involve innate antioxidant properties (Baeza, et al. 2010) or altered expression of endogenous antioxidants (Borras, et al. 2005). Indeed, estrogen protects both keratinocytes and fibroblasts from oxidative damage (Bottai, et al. 2013). Alternatively, estrogen directly regulates protease activity, inhibiting expression of MMPs −1, −2, −8, −9 and −13 (Pirila, et al. 2002; Pirila, et al. 2001; Son, et al. 2005) while promoting TIMP expression (Chen, et al. 2003). Estrogen also reduces senescence in endothelial progenitor cells by increasing telomerase levels (Imanishi, et al. 2005), and has been shown, at the level of gene expression, to confer many of the effect of ageing on delayed skin wound healing (Hardman and Ashcroft 2008).

Given the close association between estrogen level and phenotypic age-associated tissue remodelling in human skin, we hypothesised that acute estrogen deprivation (ovariectomy, Ovx) in young mice would reproduce the compositional, structural and mechanical consequences of long-term skin ageing. Using a combination of biomechanical measurements, mass spectrometry (MS) proteomics and imaging methods we compared the skin of Ovx, aged and young mice. We found that estrogen abrogation and ageing had similar effects on the ECM proteome, but that they elicited contrasting effects on skin mechanical properties, suggesting compounded mechanisms of dermal ageing.

## MATERIALS AND METHODS

### Aged and ovariectomised (Ovx) mice

Animal experiments were performed under UK Home Office licence (40/3713) following local ethics committee approval. Experimental mice were wild-type C57/Bl6 females. Mice aged 6 weeks, 12, 20 and 24 months were sourced from Charles River Laboratories (Kent, UK). Bilateral ovariectomy (Ovx) was carried out on mice aged 7 weeks as previously described (Emmerson, et al. 2012). Briefly, the mice were anaesthetised using a mixture of oxygen, nitrous oxide and isofluorane. Ovaries were exposed via ventral laparotomy, and excised using sharp sterile scissors. Incisions were closed using sutures and buprenorphine (0.1 mg/kg) administered as analgesic. Mice were sacrificed for analysis 7 weeks post-Ovx (at 14 weeks of age). All mice had age matched intact controls (*n* = 5 − 6 per group for all groups).

### Macro-mechanical testing of murine skin

Ventral skin was shaved, excised and stored in cold PBS. 10 × 40 mm strips of skin were gripped with sandpaper (5 mm at each end), clamped into an Instron 3344 100 N load cell (Instron, USA) and loaded to failure at a rate of 20 mm/min (Cooper, et al. 2015). Young’s moduli were calculated from linear sections of stress/strain curves. Tensile strength was defined as the stress at breaking. Stress-relaxation data was collected on an Instron 5943 10 N load cell (Instron), in a PBS water bath. Skin was cyclically loaded to 1 N at a rate of 10 mm/min five times to precondition the tissue, then loaded to 1 N and held at a constant strain for 160 s. Data was collected using Bluehill software (Instron).

### Quantitative mass spectrometry (MS) of skin tissue

Ventral skin was shaved and scraped to remove hair, then dissected into ~1 mm^3^ pieces and solubilised in 100 μL of 8 M urea (Fisher Scientific), or 3 M NaCl (Fisher Scientific), in 25 mM ammonium bicarbonate buffer containing 25 mM dithiothretol (DTT, Sigma), protease inhibitor cocktail (Sigma), sodium fluoride (Sigma) and sodium orthovanadate (Sigma). Six 1.6 mm steel beads (Next Advance Inc.) were added to the tube and samples were homogenised with a Bullet Blender (Next Advance Inc.) at maximum speed for 3 minutes. Resulting homogenates were cleared by centrifugation (12 °C, 10000 rpm, 5 minutes).

Immobilized-trypsin beads (Perfinity Biosciences) were suspended in 150 μL of digest buffer (1.33 mM CaCl2, Sigma, in AB (25 mM ammonium bicarbonate, Sigma)) and 50 μL of matrix extract and shaken overnight at 37 °C. The resulting digest was then reduced (addition of 4 μL × 500 mM dithiothretol, Sigma, in AB; 10 min. shaking at 60 °C) and alkylated (addition of 12 μL × 500 mM iodoacetamide, Sigma, in AB; 30 min. shaking at room temperature). Peptides were acidified by addition of 5 μL × 10% trifluoroacetic acid (Riedel-de Haën) in water, and cleaned by two-phase extraction (2 × addition of 200 μL ethyl acetate, Sigma). Peptides were desalted using POROS R3 beads (Thermo Fisher) according to the manufacturer’s protocol and lyophilized. Peptide concentrations (Direct Detect spectrophotometer, Millipore) in injection buffer (5% HPLC grade acetonitrile, Fisher Scientific, 0.1% trifluoroacetic acid in deionized water) were adjusted to 300 ng/μL. Digested samples were analysed using an UltiMate 3000 Rapid Separation liquid chromatography system (Dionex Corporation) coupled to an Orbitrap Elite (Thermo Fisher Scientific) spectrometer. Peptide mixtures were separated using a gradient from 92% A (0.1% formic acid, FA, Sigma, in deionized water) and 8% B (0.1% FA in acetonitrile) to 33% B, in 104 min at 300 nL/min, using a 75 mm × 250 μm inner diameter 1.7 μM CSH C18, analytical column (Waters). Peptides were selected for fragmentation automatically by data-dependent analysis.

### Proteomics data processing

Spectra from multiple samples were aligned using Progenesis QI (Nonlinear Dynamics) and searched using Mascot (Matrix Science UK), against the SwissProt and TREMBL mouse databases. The peptide database was modified to search for alkylated cysteine residues (monoisotopic mass change, 57.021 Da), oxidized methionine (15.995 Da), hydroxylation of asparagine, aspartic acid, proline or lysine (15.995 Da) and phosphorylation of serine, tyrosine, threonine, histidine or aspartate (79.966 Da). A maximum of 2 missed cleavages was allowed. Fold-change differences in the quantity of proteins detected in different samples were calculated by linear model fitting as has been described previously (Choi, et al. 2014; Clough, et al. 2009), using code written in Matlab (MathWorks, USA). Briefly, raw ion intensities from peptides from proteins with fewer than 3 unique peptides per protein were excluded from quantification. Remaining intensities were logged and normalized by the median intensity of ECM peptides, enabling comparison between samples and thus making quantification of ECM components an intensive measure (i.e. independent of the quantity of sample). Peptides assigned to different isoforms were collected into a single “protein group” by gene name. Only peptides observed in at least 2 samples were used in quantification. Missing values were assumed as missing due to low abundance (Goeminne, et al. 2016) and were imputed randomly at the peptide level from a normal distribution centred on the apparent limit of detection for the experiment. The limit of detection was determined by taking the mean of all minimum logged peptide intensities and down-shifting it by 1.6σ, where σ is the standard deviation of minimum logged peptide intensities. The width of this normal distribution was set to 0.3σ (Tyanova, et al. 2016). Heatmaps were created using *heatmap.2()* function from the *gplots* R package (https://cran.r-project.org/web/packages/gplots/index.html), using Wald clustering with Euclidian distances.

### Histological and immunochemical analysis of skin structure

6 μm thick histological sections were prepared from formalin-fixed, paraffin embedded tissue. Total collagen was assessed using Masson’s trichrome staining; organised fibrillar collagen was stained using Picrosirius Red (McConnell, et al. 2016); elastic fibres were stained using Gomori’s aldehyde-fuchsin (Gomori 1950). Immunohistochemistry for advanced glycation end products (AGE) was carried out with AGE antibody (Abcam, UK). Bound primary antibody was detected using Vectastain ABC kit and NovaRed detection kit (Vector Laboratories). Brightfield images were taken on a Nikon eclipse E600 microscope/SPOT camera. Picrosirius red images were captured using crossed polarised light which visualised organised fibrillar collagen (Leica DMRB) (McConnell et al. 2016). All images were analysed using ImageJ 1.46r software (National Institutes of Health, USA). To quantify elastic fibres, blinded images were thresholded and the pixels measured as a % area (Graham, et al. 2011). Collagen was measured using the colour deconvolution plugin for ImageJ. Immunohistochemistry images were analysed using Image Pro Plus software (Media Cybernetics).

### Statistical testing

MS data was compared using t-tests with Benjamini-Hochberg correction for false positives (*BHFDR*) (Benjamini and Hochberg 1995) (MatLab). Other data were compared using one way ANOVA with post hoc analysis or t-tests using Prism 7 (GraphPad Software, USA).

## RESULTS

### Ovariectomy and ageing confer opposing effects on the mechanical properties of murine skin

To investigate whether the compositional and mechanical changes induced by intrinsic ageing were caused by estrogen deprivation, we compared the mechanical properties of skin from Ovx, aged and young mice (with intact ovaries). There was a strong positive correlation (R^2^ = 0.97; *p* < 0.001) between age and skin tensile strength (stress at breaking): the breaking stress was found to be 50% greater at 24 months than at 6 weeks (Fig. 1A). Conversely, a comparison between the skin of animals 7 weeks post-Ovx and age-matched intact controls revealed a >30% reduction in tensile strength (*p* < 0.05). This trend was mirrored in the Young’s modulus (ratio of stress to strain; ‘stiffness’), which showed positive correlation with increasing age (R^2^ = 0.95; *p* < 0.05), while Ovx skin demonstrated reduced stiffness (*p* < 0.05; Fig. 1B). Finally, the skin’s viscoelastic properties were investigated by assessing the rate of stress relaxation. Again, opposing effects were observed in aged versus Ovx. Skin from 24-month animals relaxed more slowly (retained tension for longer) than young controls. By contrast, Ovx skin relaxed more quickly, indicative of the tissue being more yielding (Fig. 1C). Collectively, these unexpected data indicate that estrogen deprivation and intrinsic ageing elicit opposing effects on skin mechanical properties.

**Fig. 1.**
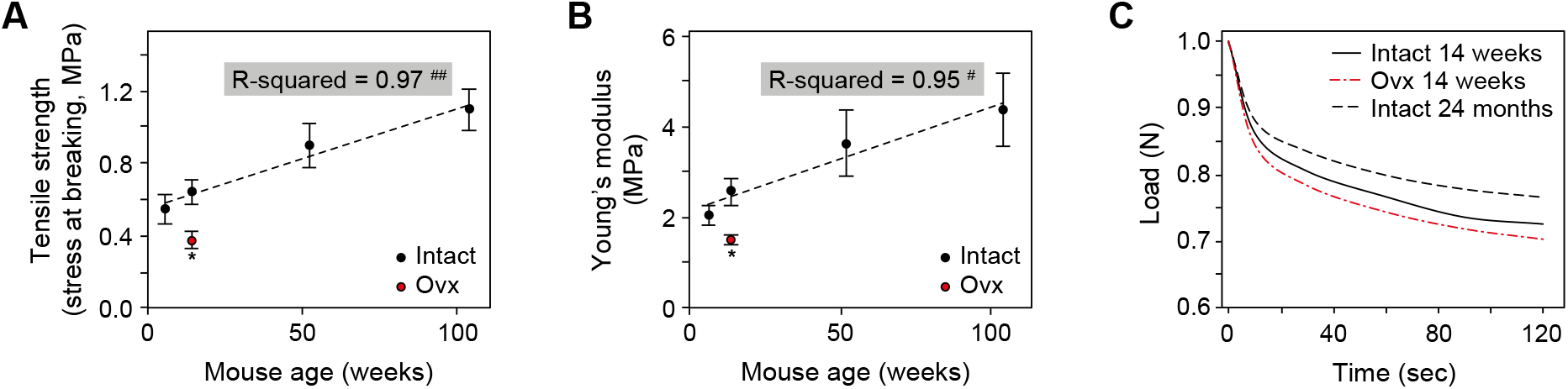
Ageing made mouse skin stronger and stiffer, ovariectomy (Ovx) had the opposite effect. (**A**) Tensile strength of mouse skin, measured as the stress at breaking, increased linearly with age. Ovx mice at 14 weeks (7 weeks following ovariectomy) had significantly reduced tensile strength, compared to age-matched controls. (**B**) Skin stiffness (Young’s modulus, ratio of stress to strain) also increased linearly with the age of the mouse. Skin stiffness of Ovx mice was significantly reduced at 14 weeks (7 weeks following ovariectomy) relative to control animals. (**C**) The viscoelastic properties of mouse skin were also altered in opposing directions in response to ageing, with stress relaxation time decreased by Ovx (7 weeks following ovariectomy) and increased by ageing (shown at 24 months) (mean ± SEM; *n* = 5 − 6; * *p* < 0.05; p-value of Pearson’s R: # *p* < 0.05, ## *p* < 0.001).

### Mass spectrometry (MS) proteomics identified characteristic changes in aged skin

To assess if changes in tissue proteome composition could be responsible for the differing tissue mechanics, we analysed mouse skin composition by MS proteomics (Fig. 2A). Ventral skin tissue was analysed from aged (20 months), young (14 weeks), and Ovx (7 weeks post ovariectomy) animals. Proteins were solubilized using either a high salt or chaotropic matrix extraction protocol and analysed by MS. The matrix proteome was defined as all proteins with ECM gene ontology annotation. Euclidean clustering revealed broad changes to the extracted proteome with animals largely grouped into the three conditions aged, young or Ovx (Fig. 2B). High salt and chaotropic extraction protocols showed complementary, overlapping ECM protein sets (Fig. 2C). For subsequent experiments, we chose chaotropic extraction due to its ability to detect elastic fibre proteins (Fbn1 and Fbn2).

**Fig. 2.**
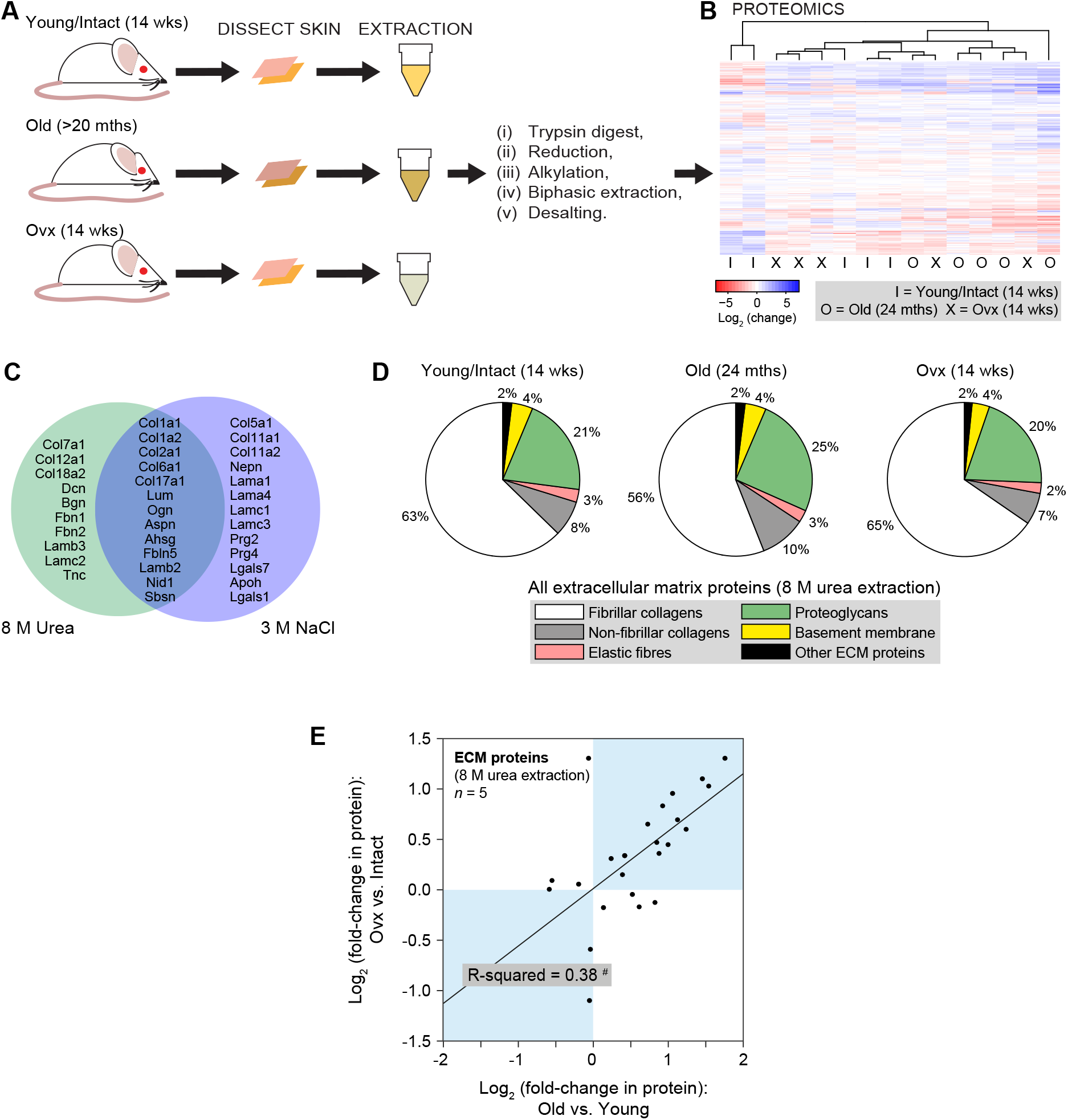
Mass spectrometry (MS) of mouse skin showed compositional changes in ageing; changes following ovariectomy (Ovx) were broadly correlated. (**A**) Schematic showing preparation and analysis of protein extracts from young, aged and ovariectomised (Ovx) mouse skin. (**B**) Heatmap showing relative protein levels (red = high; blue = low) in mouse skin extracts (I = intact/young at 14 weeks; O = old, at 24 months; X = Ovx animals at 14 weeks; *n* = 5; chaotropic extraction protocol). Columns were ordered with a Euclidean clustering algorithm, showing that proteomic features could allow separation of young and old samples while Ovx were interspersed. (**C**) Venn diagram showing proteins with ‘extracellular’ gene ontology annotation identified in mouse skin extracts using orthogonal extraction chemistries, chaotropic (8 M urea) vs. high salt (3 M NaCl). (**D**) Pie charts showing how sub-classifications of matrix proteins contributed to the total MS signal from extracellular matrix proteins, in young (14 weeks), old (24 months) and Ovx mouse skin extracts (percentages averaged *n* = 5, urea extraction protocol). (**E**) Plot showing correlation between extracellular matrix proteins (detected with three-or-more peptides per protein) extracted from skin of Ovx vs. intact and old vs. young mice (*n* = 5; significance indicated where false-discovery corrected p-value, *BHFDR* < 0.05; p-value of Pearson’s R: # *p* = 0.0013).

Next, we evaluated the proportions of the total MS signal (summed ion intensities from all derivative peptides) from ECM-derived proteins attributed to various protein sub-classifications (Fig. 2D). This comparison gives an estimate of the mass-fractions of extracted extracellular components, in contrast to measurements based on median ion intensities that estimate concentration (Ivanovska, et al. 2017). There was a relative decrease in the quantity of fibrillar collagens (Col1a1 and Col1a2) extracted from aged skin, consistent with earlier reports (Boyer, et al. 1991); instead, aged skin contained an increased proportion of signal from proteoglycans and non-fibrillar collagens.

### Comparison of ovariectomised and aged skin revealed considerable correlated changes in matrix composition

Though gross proteomic analyses revealed a greater effect of ageing than Ovx on ECM composition (Fig. 2D), analysis of changes in individual matrix proteins showed a strong correlation between Ovx and aged groups, most likely reflecting the contribution of estrogen deficiency to intrinsic ageing. Specifically, a plot of changes to ECM proteins in Ovx versus ageing (Fig. 2E) revealed correlation in 16 of 24 matrix proteins (based on proteins falling in quadrants in the *y* = *x* diagonal). A least-squares fit had a gradient of less than one, reflecting a greater perturbation caused by ageing versus ovariectomy (R-squared = 0.38; *p* = 0.0013). Outliers from the Ovx versus ageing correlation included: fibrillar collagens (Col1a1 and Col1a2) that were decreased in ageing, but unaffected by acute loss of estrogen; Col6a1, which was increased by ageing but not in Ovx; and Col6a2, which was decreased in Ovx but not by ageing.

The relative effects of Ovx and ageing were examined in different subclasses of ECM proteins (Fig. 3; significance indicated where the false-discovery corrected p-value, *BHFDR* < 0.05). Multiple fibrillar collagens were significantly perturbed with ageing (Col1a1, Col1a2, Col7a1 and Col17a1), but were unaffected in Ovx (Fig. 3A). Non-fibrillar collagens – particularly type-VI – were differentially altered by ageing and Ovx (Fig. 3B). Elastic fibre proteins displayed a non-significant trend toward perturbation only in Ovx mice (Fig. 3C). Changes to levels of proteoglycan (Fig. 3D) and basement membrane proteins (Fig. 3E) were broadly correlated in Ovx versus ageing, with increased levels seen across both subclasses of protein. These results highlight the specific role(s) estrogen deficiency may play in age-related changes in tissue mechanics, and strongly suggest that estrogen regulates the levels of basement membrane proteins and proteoglycans during ageing. Furthermore, the contrasting effects of ageing and Ovx on fibrillar collagens and elastic fibres, respectively, suggest that estrogen deprivation is not a primary cause of reduced type-I collagen and that ageing may compensate for estrogen deprivation-driven loss of elastic fibres.

**Fig. 3.**
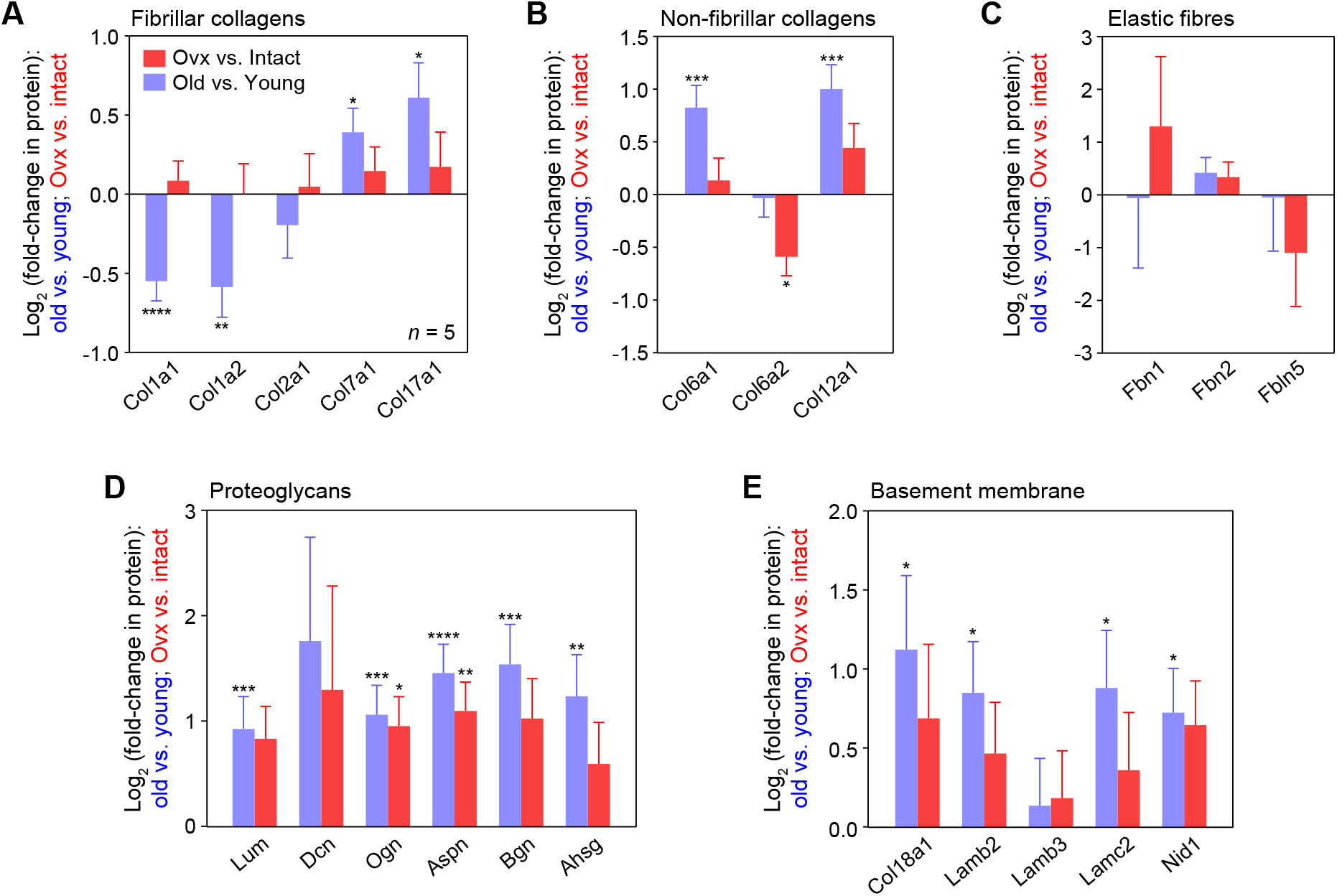
Quantitative proteomic analyses of extracellular matrix protein classes showed differential changes to mouse skin in ageing vs. ovariectomy (Ovx). Plots of extracellular matrix quantified in dermal extracts (urea protocol), comparing protein fold-changes in ageing (24 months vs. 14 weeks; *n* = 5 animals) and acute estrogen loss (Ovx, 7 weeks after ovariectomy, normalized to intact control, both at 14 weeks; *n* = 5 animals). (**A**) Fibrillar collagens; (**B**) Non-fibrillar collagens; (**C**) Elastic fibres; (**D**) Proteoglycans; (**E**) Basement membrane proteins (mean ± SEM of coefficient determined from linear modelling; false-discovery corrected significance indicated on plots: * *BHFDR* < 0.05, ** *BHFDR* < 0.01, *** *BHFDR* < 0.001, **** *BHFDR* < 0.0001).

### Histological analysis showed neither ageing nor ovariectomy changed dermal collagen density or organization

As collagens are abundant in dermal tissue, important in determining mechanical properties (Swift, et al. 2013) and were specifically down-regulated during ageing but not Ovx, we resolved to further characterize the organization of fibrillar proteins. Masson’s trichrome staining for total collagen revealed no observable differences between dermal tissue at 14 weeks, 20 months or 7 weeks post-Ovx (Fig. 4A). Quantification of the stain area showed no significant differences between the three samples (Fig. 4B) – note that this was evaluated as a percentage of total stain area, an intensive measure, and so independent of dermal thickness. A similar analysis of dermal collagen density also showed no individual significant changes (Fig. 4C). There was no correlation between tensile strength (Fig. 4D) or Young’s modulus (Fig. 4E) and the collagen densities of individual skin samples. Plane-polarised imaging of Picrosirius Red stained samples, which specifically identifies organised fibrillar collagen (McConnell et al. 2016), showed no notable differences (Fig. 4F). Image analysis confirmed no significant change in the density of thin or thick collagen fibres (Fig. 4G). Finally, analysis of collagen fibre orientation (Fig. 4H) revealed no significant changes to the coherence of fibre orientation with ageing or ovariectomy (Fig. 4I).

**Fig. 4.**
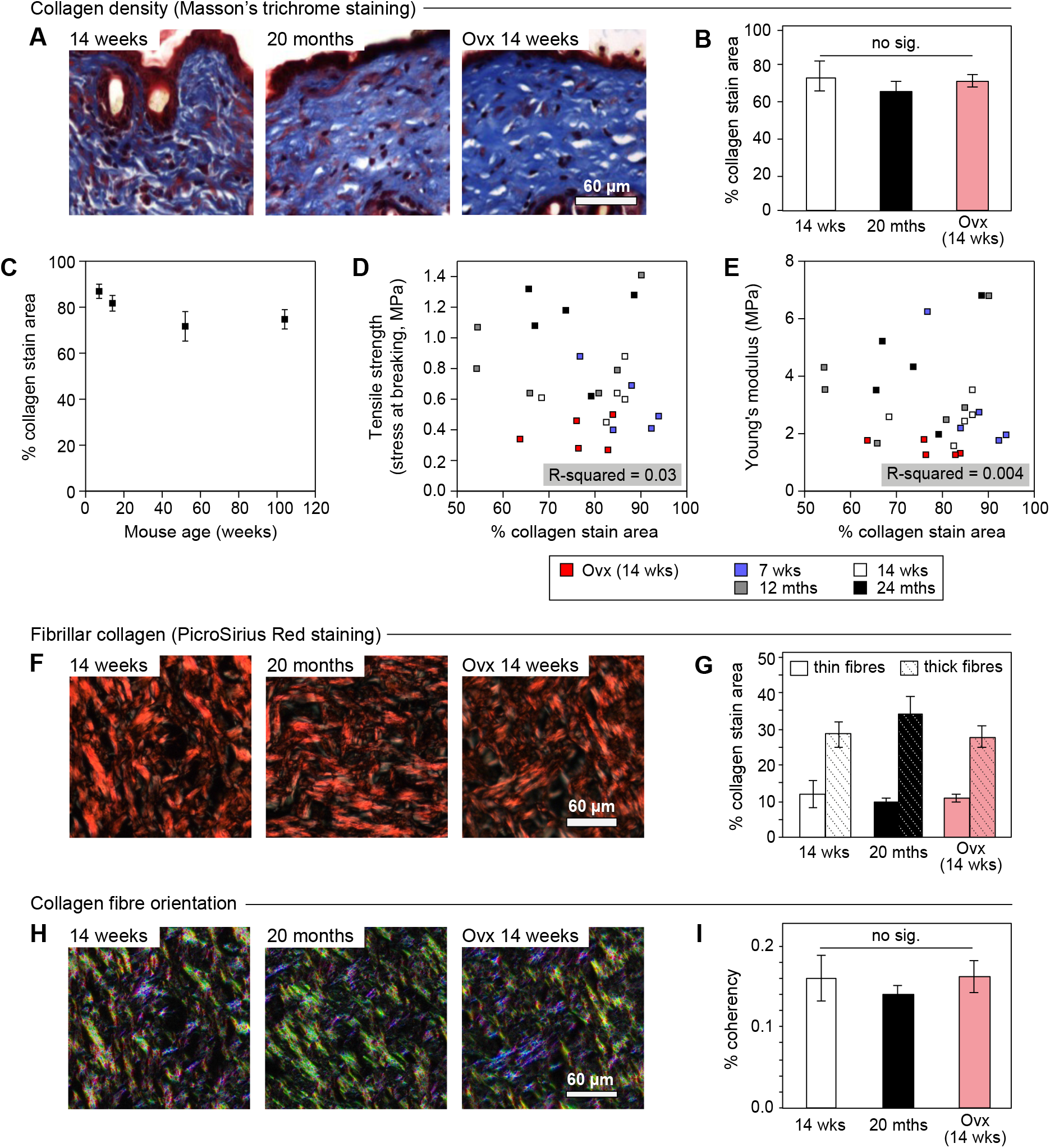
Ageing and ovariectomy (Ovx) did not affect collagen density, fibre formation or orientation in murine dermal tissue. (**A**) Total dermal collagen stained by Masson’s trichrome. (**B**) Quantification of dermal collagen density showed no significant change with animal age (20 months versus 14 weeks) or following Ovx (at 14 weeks, 7 weeks after ovariectomy). (**C**) Dermal collagen density measured at 7 and 14 weeks, and at 12 and 24 months; there were no significant differences between any of the individual age groups. (**D**) Dermal collagen density did not correlate with the tensile strength of the sample (measured as the stress at breaking; R-squared = 0.03). (**E**) There was also no correlation between dermal collagen density and the Young’s modulus of the sample (R-squared = 0.004). (**F**) Fibrillar collagen imaged with Picrosirius Red staining (McConnell et al. 2016). (**G**) Quantification of Picrosirius Red staining against fibrillar collagen showed that neither Ovx or ageing significantly affected the density of thin or thick fibrils. (**H**) Collagen staining with polarization to contrast fibre orientation. (**I**) Quantification of collagen fibre orientation showed no significant variation with Ovx or increasing age (mean ± SEM; *n* ≥ 5).

### Elastic fibres were decreased following ovariectomy, while advanced glycation end-products (AGEs) increased with ageing

Histological analysis of skin elastic fibres using Gomori’s aldehyde-fuchsin staining (Fig. 5A) revealed a statistically significant 39% decrease in elastic fibre content in the skin of ovariectomised mice (7 weeks post-Ovx; *p* < 0.001), with no significant change in aged mouse skin (Fig. 5B). Immunohistochemical localisation of the dermal elastic fibre component protein fibrillin-1 (Fig. 5C), revealed a similar profile: a significant reduction in Ovx skin, with no significant change in aged skin (*p* < 0.05; Fig. 5D). Plots of tensile strength (Fig. 5E) and Young’s modulus (Fig. 5F) versus fibrillin-1 abundance showed power law scaling when considering data from 7-week, 14-week and Ovx samples (*p* < 0.05). Similar scaling relationships have been reported in studies of the dependence of elasticity on the concentration of biological polymers (Gardel, et al. 2004; Swift et al. 2013). However, scaling was lost in 12-month and 24-month samples, where tensile strength and Young’s modulus were increased without a corresponding increase in the abundance of fibrillin-1. This result suggests that additional, non-estrogenic, mechanisms of tissue stiffening emerge in ageing.

**Fig. 5.**
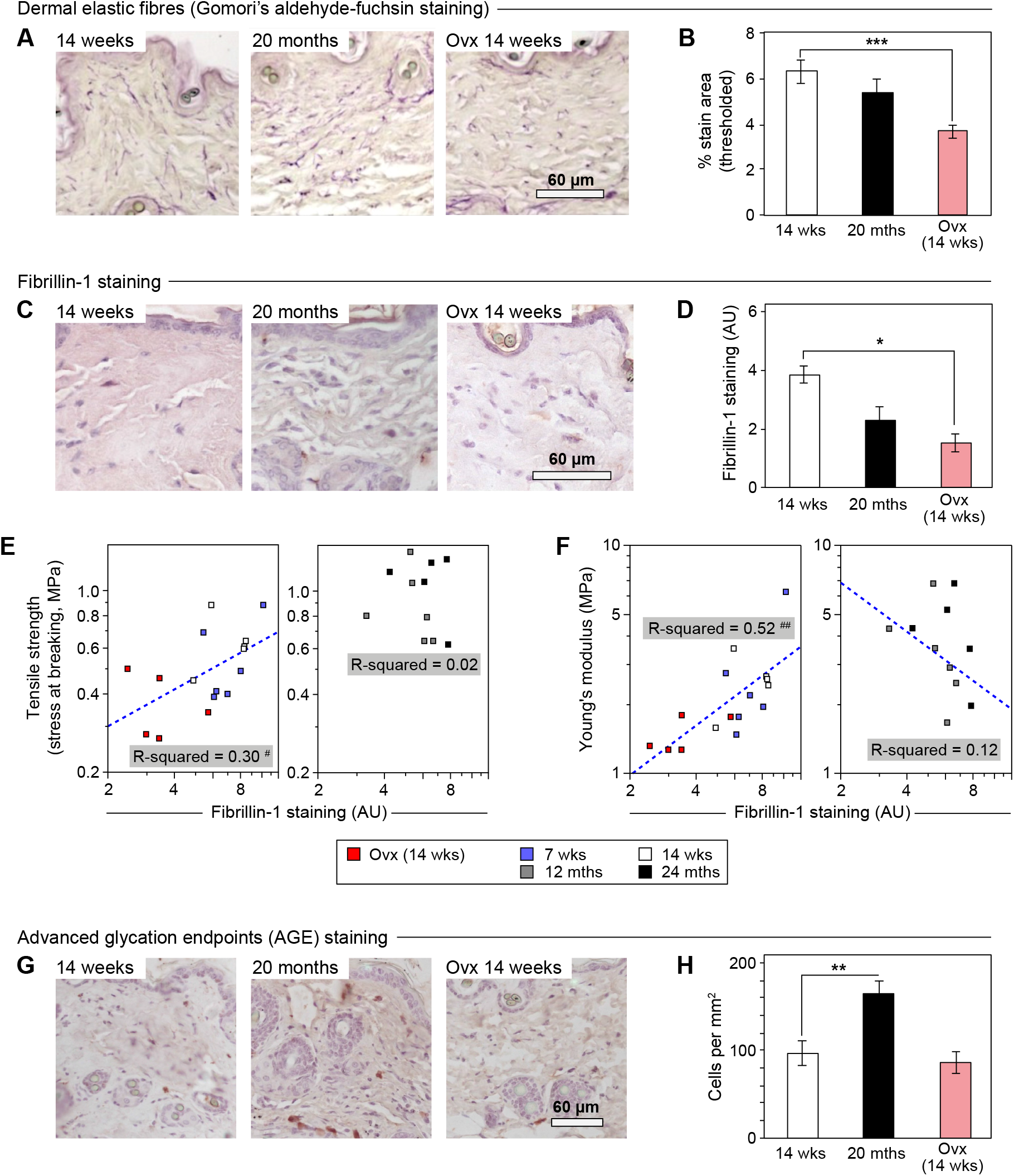
Ovariectomy (Ovx) decreased density of elastic fibres in murine dermal tissue; advanced glycation endpoints (AGE) increased with age. (**A**) Dermal elastic fibres visualized by Gomori’s aldehyde-fuchsin staining (Gomori 1950). (**B**) Quantification of staining showed that dermal elastic fibre density was reduced by ageing (14% decrease at 20 months) and by Ovx at 14 weeks (7 weeks after ovariectomy; 42% decrease), relative to 14 week-old control animals. (**C**) Immunohistochemical staining against elastic fibre component fibrillin-1. (**D**) Quantification of fibrillin-1 staining showed a corresponding, significant reduction following Ovx. (**E**) Plots of tensile strength versus fibrillin-1 staining intensity for discrete skin samples. Young mouse skin (Ovx 14 weeks, intact 7 and 14 weeks) showed positive scaling (R-squared of combined data = 0.30), although this was not observed in older skin samples (12 and 24 months). (**F**) Plots of Young’s modulus versus fibrillin-1 intensity for discrete samples. Young tissue (Ovx 14 weeks, intact 7 and 14 weeks) showed positive power-law scaling (R-squared of combined data = 0.52); older skin samples (12 and 24 months) did not show the same scaling. (**G**) Immunohistochemical staining against advanced glycation endpoints (AGE). (**H**) Quantification of AGE staining showed a significant increase in the density of positively stained cells at 20 months compared with 14 weeks. Figure parts (B), (D) and (H): mean ± SEM; *n* ≥ 5; * *p* < 0.05, ** *p* < 0.005, *** *p* < 0.001. Figure parts (E) and (F): p-values of Pearson’s R: # *p* < 0.05, ## *p* < 0.005).

The MS quantified decrease in type-I collagen (Fig. 3) with age seemed counter-intuitive to the observed increased skin stiffness (Fig. 1) and the minor changes to collagen-I staining (Fig. 4). We hypothesised that these results, along with the lack of dependence of stiffness on fibrillin-1 in ageing skin (Fig. 5), could be explained by age-induced crosslinking of ECM proteins. Reactive intermediates of glycolysis can accumulate with age and promote non-enzymatic crosslinking through the formation of advanced glycation end-products (AGEs). ECM proteins that can be subject to glycation and subsequent crosslinking include type-I collagens and elastin (Kasper and Funk 2001). Interestingly, highly crosslinked protein networks are stiffer (Gardel et al. 2004; Kim, et al. 2009), and less soluble and thus would be detected at lower concentrations by MS. Histological quantification of AGEs (Fig. 5G) revealed a significant increase at 20 months versus 14 weeks (*p* < 0.005), but no change post-Ovx (Fig. 5H). Thus, increased AGE-mediated crosslinking in ageing skin could underlie the increased stiffness that is not observed in ovariectomised mice, despite similarities observed by proteomic analysis.

## DISCUSSION

### Comparison of mechanical changes in ageing and following ovariectomy

The influence of age-related hormonal changes on skin mechanobiology is poorly understood. In this study we present novel data demonstrating that ageing and estrogen deficiency exert opposing effects on the mechanical properties of mouse skin. Ageing increased skin tensile strength and stiffness, as well as altering viscoelastic properties. By contrast, a short (7 week) period of ovariectomy caused skin tissue to be weaker, more compliant and to relax more quickly following stress. Our finding that the Young’s modulus of mouse skin increases with age is in line with the widely accepted view that ageing leads to stiffening in a range of tissues, such as lung (Calhoun, et al. 2016) and muscle (Phillip, et al. 2015). Indeed, skin could provide a readily accessible model to study conserved features of the mechanobiology of ageing elastic tissues. Curiously, the reported effects of ageing on human skin are less clear: torsion testing *in vivo* has shown the Young’s modulus to be increased by ageing (Agache, et al. 1980) while dynamic indentation measurements have suggested the opposite result (Boyer, et al. 2009). This variation likely reflects the inherent difficulties in performing consistent measurements on living human tissue (reviewed in Moronkeji and Akhtar 2015). The very short time frame in which estrogen deficient (Ovx) skin became weaker and more compliant is fascinating, given that it mirrors the phenotype of advanced human ageing where (particularly female) skin becomes lax, fragile and more susceptible to skin tears (El-Domyati, et al. 2002; Kennedy and Kerse 2011).

### Altered mechanical properties in aged skin may be due to crosslinking

The mechanical properties of tissue can be broadly attributed to fibrillar collagen content, where higher concentrations increasingly drive stiffening (Swift et al. 2013). However, we detected less fibrillar collagens by MS in aged skin. There are two possible interpretations for this observation. Firstly, that there is less collagen in the aged murine skin, consistent with reports of decreased collagen-I transcription with age in human skin (Makrantonaki, et al. 2012). However, this is unlikely given the typically long half-life of collagen-I molecules. Note, histologically we observed only a small (~10% over two years), non-significant reduction in collagen staining. This leads to a second interpretation, that changes to post-translational modification (PTM) states of collagen in aged tissue – in particular intramolecular crosslinking – may render tissue more resistant to collagen extraction.

In line with this second interpretation, we observed increased levels of advanced glycation end products (AGEs) in aged murine tissue, in agreement with studies reporting increased AGE-induced collagen crosslinking in aged human skin (Fan, et al. 2010). This could explain the disparity in our study between a) collagen measured by MS and collagen measured by histology and b) tissue collagen levels and tissue mechanics. PTM may in fact be a major contributor to age-associated alterations in mechanical properties, as AGEs can contribute to crosslinking of collagen and thus lead to increased stiffness (Verzijl, et al. 2002). Further, the high lysine content and relatively low turnover rate makes collagen accumulation of AGE-induced crosslinks more likely (Verzijl, et al. 2000). Note, prior studies have directly linked AGE crosslinking to altered mechanical properties in tendon (Gautieri, et al. 2017; Reddy 2004). It is important to note that our data (Ovx for 7 weeks) do not rule out a role for estrogen deficiency in long-term AGE accumulation (over the order of months and years).

### Common compositional features in ECM of ovariectomised and aged mouse skin

MS analysis of whole skin samples from ovariectomised versus intact, and aged versus young mice, revealed a high level of correlation, particularly in proteoglycans and basement membrane proteins. Although we noticed differences in levels of the highly abundant type-I collagens only in aged skin, analysis of the wider ECM proteome indicated correlated changes between ageing and ovariectomy in protein groups, particularly in proteoglycans and basement membrane proteins. Both ovariectomy and ageing increased levels of key small leucine-rich repeat (SLRP) (lumican, decorin, asporin, mimecan and biglycan) and basement membrane proteins. The SLRPs play important roles in collagen fibrillogenesis, controlling the correct formation and spacing of the collagen fibres, which will in turn alter the mechanical properties of the tissue. *In vitro* lumican and decorin knockout mice show skin fragility due to alterations in the spacing, size and formation of collagen fibrils (Chakravarti, et al. 1998; Danielson, et al. 1997). SLRPs additionally protect collagen fibrils from proteolysis (Geng, et al. 2006), and may therefore modulate collagen turnover, and ultimately the accumulation of AGE-linked molecules. Indeed, in the tendon decorin is an essential mediator of age-related changes in both structure and mechanical properties (Dunkman, et al. 2014), while in the skin estrogen receptor (ERβ) null mice display reduced decorin levels and altered collagen architecture (Markiewicz, et al. 2013).

### Estrogen deficiency selectively impacts the skin elastic fibre network

Histological analysis showed that the elastic fibre content of mouse skin was significantly, and selectively, decreased following ovariectomy. Elastic fibres principally comprise an elastin-rich core surrounded by an outer mantle of fibrillin microfibrils. The elastin core itself is highly crosslinked and so difficult to extract from tissue and subsequently quantify by MS. However, we were able to detect fibrillins (Fbn1 and Fbn2) and the fibril-associated fibulin (Fbn5). Fbn5 is essential for formation, arrangement and stabilisation of new elastic fibres and mutations are known to cause the skin laxity disease, cutis laxia (Berk, et al. 2012), and other disorders caused by ECM dysfunction. Thus, reduced Fbn5 in estrogen-deprived skin may underpin the greater elastic fibre changes versus aged tissue. Ageing and ovariectomy also conferred contrasting effects on type-IV collagen, with ageing increasing levels of Col6a1 and ovariectomy decreasing Col6a2. Although collagen-VI mutations have been characterised in muscular dystrophies (in many cases presenting with a roughened skin phenotype; Bonnemann 2011), and the importance of collagen-VI in the cancer microenvironment has been recognised (Chen, et al. 2013), relatively little is known about roles of type-VI collagens in mechanobiology and ageing.

### Quantification of ECM composition in whole tissues by MS

Extraction and solubilisation methods that are effective for analysis of the intracellular proteome (e.g. buffers containing surfactants such as sodium dodecyl sulphate, SDS) are often less effective at extracting ECM. We therefore trialled two alternative conditions to extract ECM proteins from mouse skin (Barallobre-Barreiro, et al. 2016). While ionic disruption using high salt extracted more unique ECM proteins in total, we used chaotropic disruption with urea as this yielded elastic fibre proteins (fibrillin and fibulin) that were absent from the high salt extraction method. ECM proteins are subject to a range of PTMs, including crosslinking, that can complicate detection by MS. Here, disulphide crosslinks were removed during sample preparation by performing ECM extractions under reducing conditions. Hydroxylated peptides were detected in MS by enabling identification of proline and lysine residues with masses increased by the addition of an oxygen atom. Glycosylation results in the addition of highly variable sugar chains to specific residues. The variable nature of this PTM, and the resulting variety of masses it can add to peptides, means that specific strategies are needed to analyse glycosylation patterns by MS (Ahn, et al. 2015). We therefore excluded glycosylation-associated mass changes from variable modification Mascot searches in this study. Nonetheless, glycosylation patterns have been shown to change with age and have been implicated in neurodegenerative and metabolic disease, as well as general frailty (Miura and Endo 2016).

## Conclusion

Changes in the mechanical properties of tissues are associated with a variety of diseases, along with general frailty and morbidity. In this study, we used the ovariectomised mouse model to explore the contribution of estrogen–deprivation to the changes in dermal composition and skin mechanics with age. Although ovariectomy-induced changes to the ECM proteome largely correlate with those observed in ageing, diametrically opposed changes in tissue mechanics were observed. Aged mouse skin was found to be stronger and stiffer, which we primarily attribute to protein PTMs, such as the formation of crosslinks between collagen molecules. By contrast, ovariectomy specifically altered skin elastic fibres, leading to increased yielding and reduced breaking stress. A summary of observations of mechanical and compositional properties of aged and ovariectomised mouse skin can be found in Table 1. An important finding from this study was the short time-frame over which estrogen deprivation led to significant effects on tissue mechanics and the ECM proteome, i.e. just seven weeks after ovariectomy. This implies that prompt intervention at the time of menopause to maintain both protein homeostasis and tissue mechanical properties could have significant therapeutic benefit. As the population continues to age it is important to understand how hormone deficiency interacts with chronological ageing to alter structural and mechanical features of tissues. This in turn should highlight opportunities for therapeutic intervention, such as by targeting pathways of mechano-signalling, or by directly modifying the tissue mechanical properties (Mallikarjun and Swift 2016).

**Table 1.** Summary of observations in mouse skin, comparing the effects of ageing and ovariectomy (Ovx).

|  | Old vs. Young<br>(at > 20 months) | Ovariectomy (Ovx) vs. Intact<br>(14 weeks, 7 weeks post-ovariectomy) |
| --- | --- | --- |
| **Tensile strength**<br>(strain at breaking) | ↑ | ↓ |
| **Stiffness**<br>(Young's modulus) | ↑ | ↓ |
| **Stress relaxation time**<br>(viscoelastic response) | ↑ | ↓ |
| **Extractable type-I collagen**<br>(mass spectrometry) | ↓ | No change |
| **Extractable type-VI collagen**<br>(mass spectrometry) | ↑ (Col6a1) | ↓ (Col6a2) |
| **Collagen density & alignment**<br>(histology and staining) | No change | No change |
| **Extractable elastic fibre proteins**<br>(mass spectrometry) | No change | No change |
| **Elastic fibres**<br>(histology and staining) | No change | ↓ |
| **Extractable proteoglycans**<br>(mass spectrometry) | ↑ | ↑ (Ogn, Aspn) |
| **Advanced glycation endpoints**<br>(AGE; staining) | ↑ | No change |

## SUPPLEMENTARY DATA

Proteomics data have been deposited to the ProteomeXchange Consortium via the PRIDE partner repository (Vizcaino, et al. 2013) with the identifiers PXD012753 and PXD012754.

## DECLARATION OF INTEREST

The authors declare no competing or financial interests.

## FUNDING

This work was supported by an AgeUK Senior Fellowship to MJH and a Biotechnology and Biological Sciences Research Council (BBSRC) David Phillips Fellowship (BB/L024551/1) to JS. VM was supported by the Sir Richard Stapley Educational Trust. The Wellcome Centre for Cell-Matrix Research, University of Manchester, was supported by core funding from the Wellcome Trust (203128/Z/16/Z).

## AUTHOR CONTRIBUTIONS

CRS, MJH and MJS conceived the study; CRS performed all experiments unless otherwise indicated, analysed the data, wrote and co-edited the manuscript; VM and JS performed mass spectrometry analysis; EE and DFH performed mechanical characterisation; BD, VM, JS, MJS and MJH co-edited the manuscript.

## ACKNOWLEDGEMENTS

Proteomics analysis was carried out at the Wellcome Centre for Cell-Matrix Research Biological Mass Spectrometry Core Facility. We thank Drs Ronan O’Cualain and David Knight for proteomics support.

